# ASM-Clust: classifying functionally diverse protein families using alignment score matrices

**DOI:** 10.1101/792739

**Authors:** Daan R. Speth, Victoria J. Orphan

## Abstract

Rapid advances in sequencing technology have resulted in the availability of genomes from organisms across the tree of life. Accurately interpreting the function of proteins in these genomes is a major challenge, as annotation transfer based on homology frequently results in misannotation and error propagation. This challenge is especially pressing for organisms whose genomes are directly obtained from environmental samples, as interpretation of their physiology and ecology is often based solely on the genome sequence. For complex protein (super)families containing a large number of sequences, classification can be used to determine whether annotation transfer is appropriate, or whether experimental evidence for function is lacking. Here we present a novel computational approach for de novo classification of large protein (super)families, based on clustering an alignment score matrix obtained by aligning all sequences in the family to a small subset of the data. We evaluate our approach on the enolase family in the Structure Function Linkage Database.

**Availability and implementation:** ASM-Clust is implemented in bash with helper scripts in perl. Scripts comprising ASM-Clust are available for download from https://github.com/dspeth/bioinfo_scripts/tree/master/ASM_clust/

## Introduction

The rapid advances in sequencing technology have led to a dramatic increase in available genome sequences. This genomic data has provided new perspectives on big questions in biology, such as the diversity of life, the distribution of metabolic traits across the tree of life, and the origin of eukaryotes (Hug et al. 2016; Zaremba-Niedzwiedzka et al. 2017; Borrel et al. 2019). In addition, each newly available genome sequence contains novel protein sequences, yielding novel protein families of unknown function and expanding families with previously characterized representatives. Automatic functional annotation of novel protein sequences is generally done by annotation transfer from known homologous proteins, either using sequence alignment or hidden markov models (Altschul et al. 1990; Finn, Clements, and Eddy 2011). This approach can, and often does, result in misinterpretation of the function of proteins in mechanistically diverse superfamilies, and is prone to subsequent error propagation (Schnoes et al. 2009). Accurately interpreting the function of novel proteins is one of the grand challenges in biology, and relies heavily on availability of experimental data. Classifying mechanistically diverse protein superfamilies provides insight in knowledge gaps, can indicate whether annotation transfer is appropriate, and can help guide future experiments.

There are various automatic tools available for classification of proteins into isofunctional families using sequence similarity, active site characteristics, and phylogenetic relationships (Brown, Krishnamurthy, and Sjölander 2007; Lee, Rentzsch, and Orengo 2010; de Melo-Minardi, Bastard, and Artiguenave 2010; Leuthaeuser et al. 2016; Knutson et al. 2017). Alternatively, the structure of a protein family can be interactively explored using sequence similarity networks (SSNs) (Atkinson et al. 2009; Copp et al. 2018). SSNs are constructed based on pairwise all vs all alignment, with each node in the network representing a sequence, and each edge between two nodes representing the alignment between sequences. Clusters of nodes can be manually selected, or identified using a clustering algorithm such as MCL (Enright, Van Dongen, and Ouzounis 2002). SSNs are a powerful method to investigate protein families, but the network visualization limits the number of sequences that can be included, and alignments between separate domains of multi-domain proteins may confuse the analysis.

Here we present ASM-Clust, an alternative method that uses alignment score matrix (ASM) clustering. For each input sequence, ASM-Clust generates a profile consisting of a large number of alignment scores, including both presence/absence and weight, and uses this profile to classify each sequence. For a dataset containing *N* sequences, alignments are generated for all N sequences against a randomly selected subset of *n* sequences, and taking each alignment score, or a zero if the sequence did not align to the reference. This results in a matrix of *N* x *n* values which is subsequently visualized using t-distributed stochastic neighbor embedding (t-SNE) (Van der Maaten and Hinton 2008; Van der Maaten 2014), and can be clustered using DBscan (Ester et al. 1996).

## Implementation

ASM-Clust is implemented in bash with helper scripts in perl, and will take a protein fasta file as the sole input. Fasta files are processed with ASM_clust.sh, which 1) randomly selects a subset of n sequences (default 1000), 2) aligns the entire dataset to the subset of n sequences, 3) combines all scores into a matrix (inserting 0 for query-database pairs that did not produce an alignment), and 4) reduces the matrix to 2 dimensions using t-SNE (Figure 1a). For flexible usage, ASM-Clust supports alignment using DIAMOND (Buchfink, Xie, and Huson 2015), BLAST (Altschul et al. 1990), or MMSeqs2 (Steinegger and Söding 2017), and uses MMSeqs2 as default alignment software. Clustering results are comparable between different alignment software (Supplemental Figure S1). Other user-defined options are the number of sequences in the subset (default 1000), the main t-SNE parameter “perplexity” (default 1000) and maximum iterations (default 5000) for dimensionality reduction, and the number of threads used by the alignment software (default 1). Although the clustering is generally similar with multiple randomly chosen subsets (Supplemental Figure S2), the subset can be defined for reproducibility. The output of ASM_clust.sh can be visualized as a scatterplot where each dot represents a sequence, and clusters are readily apparent (Figure 1b, Supplemental figure S1-S3). This format allows additional annotation with sequence features, such as taxonomy, length, or composition. The visualization in Figure 1b and Supplementary Figures S1-S3 was created using R, with the ggplot2 package, and clusters were called using the dbscan package. The t-SNE result and the annotation data downloaded from SFLD were combined prior to visualization. Clusters obtained with ASM-Clust can be further refined by iteratively applying the method to a subset of poorly resolved data, such as the “hub”s cluster (Supplementary Figure S3). The smaller total number of sequences in the second iteration, combined with lowering the perplexity value of the t-SNE, increases the resolving power of the analysis for clades with a small number of sequences, thus resolving rare classes with few members (Supplementary Figure S3).

**Figure 1.**
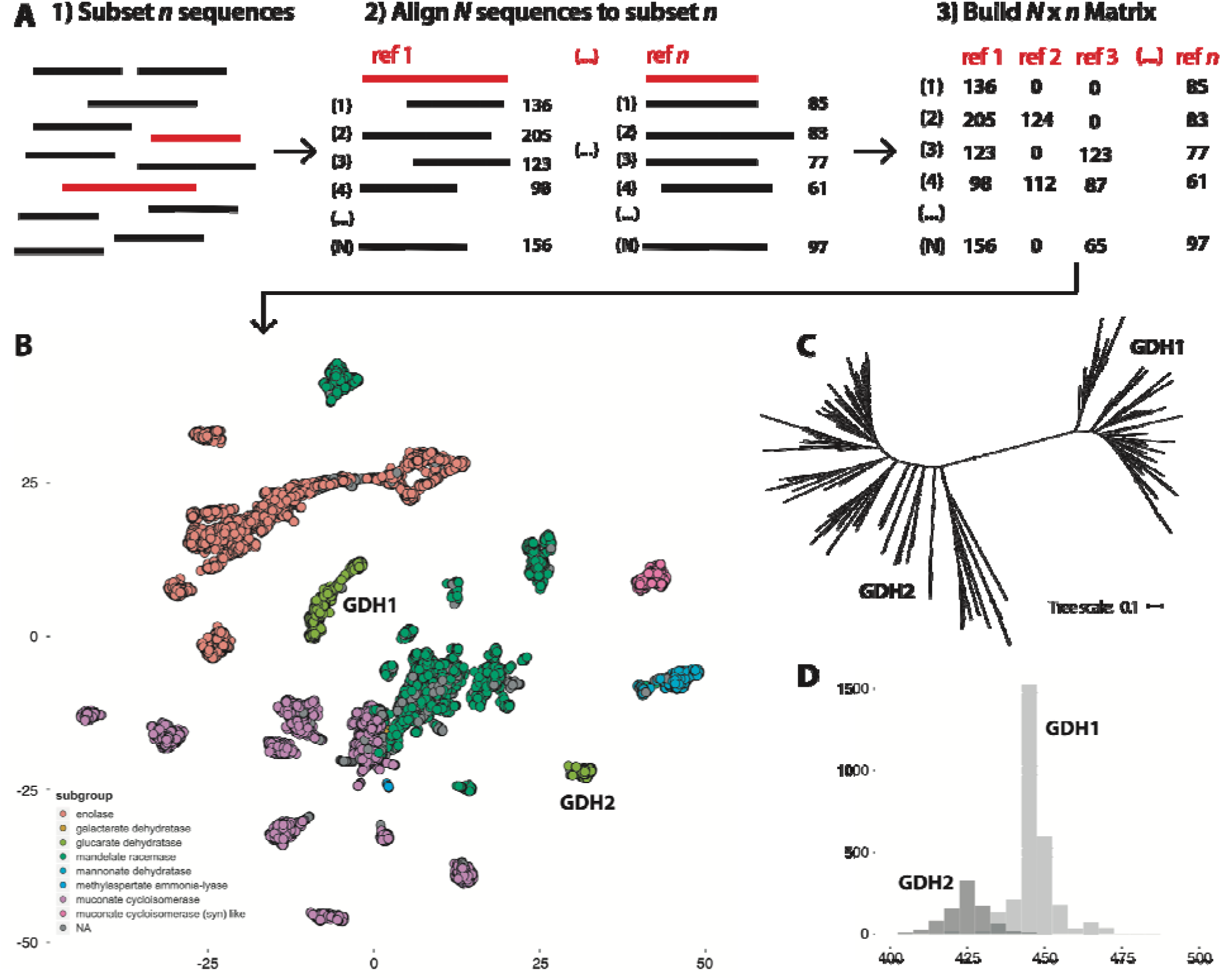
ASM-clust workflow overview and enolase superfamily example. A) ASM-Clust workflow overview and B) example of the ASM-Clust output on the structure function linkage database (SFLD) enolase superfamily (48,850 sequences). The clusters are colored by SFLD subgroup: enolase (red), galactarate dehydratase (orange), glucarate dehydratase (light green), mandelate racemase (dark green), mannonate dehydratase (light blue), methylaspartate-ammonia lyase (dark blue), muconate cycloisomerase (purple), muconate cycloisomerase (syn) like (pink), and no assigned subgroup (gray). The isofunctional ‘glucarat dehydratase’ subgroup is split in two clusters, indicated with GDH1 and GDH2. C) Phylogenetic analysis of the ‘glucarate dehydratase’ subgroup (clustered at 70% identity), and D) sequence length comparison confirms the clear separation of the two clusters.

## Results

ASM-Clust was tested on the enolase superfamily in the gold-standard Structure Function Linkage Database (SFLD) (Akiva et al. 2014). Sequences and annotation data table were downloaded from the SFLD website (http://sfld.rbvi.ucsf.edu) and all 48,850 sequences were clustered using ASM-Clust with default settings, and visualized using R (Fig 1b). The ‘mannonate dehydratase’ and ‘muconate cycloisomerase (syn) like’ subgroups, each containing only a single isofunctional family, are well resolved. As expected, the functionally diverse ‘muconate cycloisomerase’ and ‘mandelate racemase’ subgroups each partition into multiple discrete clusters (Figure 1b). The isofunctional ‘enolase’ and ‘glucarate dehydratase’ subgroups also result in multiple clusters (Figure 1b). Phylogenetic analysis of the ‘glucarate dehydratase’ subgroup confirms that the observed clusters respond to distinct clades that can also be separated by sequence length (Figure 1c & 1d, Supplementary methods). The smaller methylaspartate ammonia-lyase and galactarate dehydratase subgroups (307 and 25 sequences respectively) are more clearly resolved when ASM-clust is iteratively rerun on the “hub” cluster with a lower perplexity value (Supplemental Figure S3). When prior high-quality annotation is not available, clusters can be inspected for phylogeny, taxonomic distribution, and conserved residues to assess whether they represent functionally divergent sequences.

ASM-Clust can retrieve clades from a complex superfamily with tens of thousands of sequences, without prior reduction of the dataset. We expect this to become increasingly relevant as the amount of sequence data from phylogenetically diverse organisms continues to grow rapidly, and meaningful information can be overlooked while pre-clustering a sequence dataset.

