## Supplementary material for "ASM-Clust: classifying functionally diverse protein families using alignment score matrices": Suppl Methods and Suppl Figs S1-S3

After visualization of the t-SNE result for the enolase superfamily, clusters were assigned using the dbscan package, with epsilon neighbourhood of 3. Clusters were exported as tab delimited files, and sequences of the two glucarate dehydrogenase clusters were obtained using the tab_seq_lookup.pl script (https://github.com/dspeth/bioinfo_scripts/tree/master/ASM_clust/). To reduce the number of sequences prior to phylogenetic analysis either set was clustered at 70% identity using UCLUST (Edgar, 2010) yielding 56 and 90 sequences for GDH1 and GDH2 respectively. Clustered sequence sets were combined and the resulting 146 sequences were aligned using MUSCLE (Edgar, 2004) and a maximum likelihood phylogeny was calculated using RAxML (Stamatakis, 2014), with the LG4X substitution model (Le, Dang & Gascuel, 2012) and 500 bootstrap replicates. The phylogeny was visualized using iTOL (Letunic & Bork, 2016).

Protein length was calculated using the fasta_to_tab_id_length.pl script (https://github.com/dspeth/bioinfo_scripts/blob/master/proteins/), and vizualized in R using the ggplot2 package.

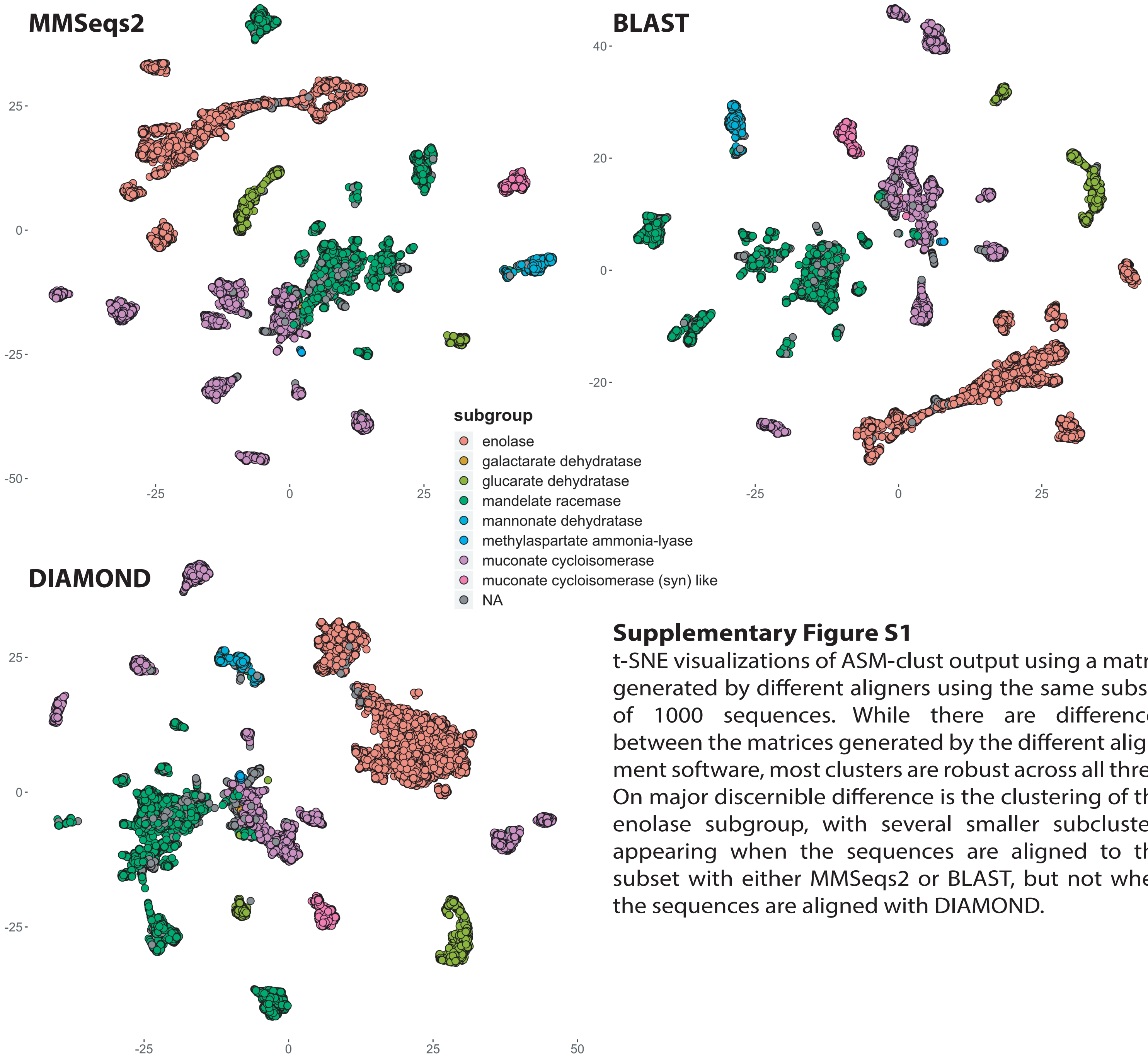


**Supplementary Figure S1**

t-SNE visualizations of ASM-clust output using a matrix generated by different aligners using the same subset of 1000 sequences. While there are differences between the matrices generated by the different alignment software, most clusters are robust across all three. On major discernible difference is the clustering of the enolase subgroup, with several smaller subclusters appearing when the sequences are aligned to the subset with either MMSeqs2 or BLAST, but not when the sequences are aligned with DIAMOND.

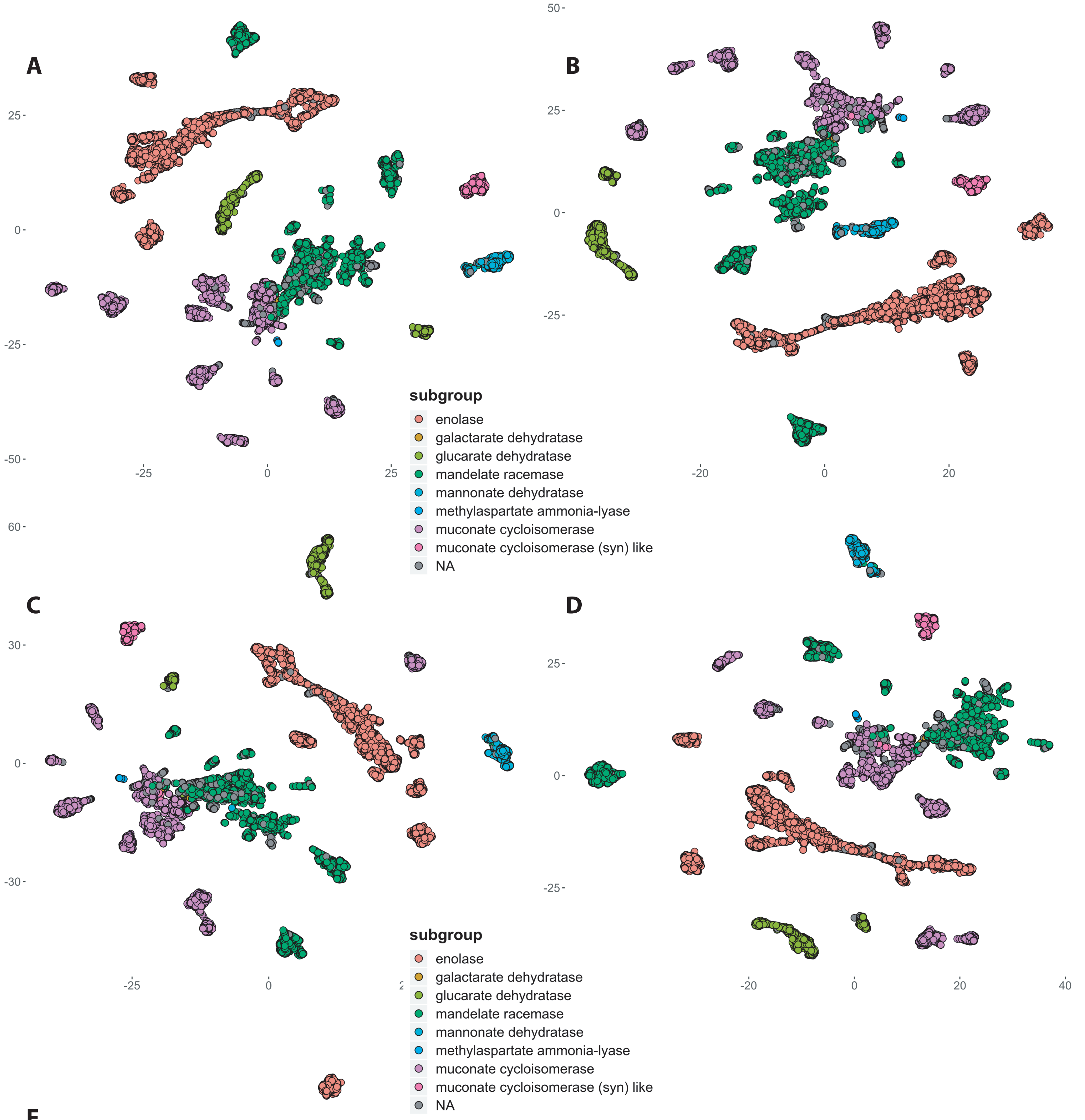


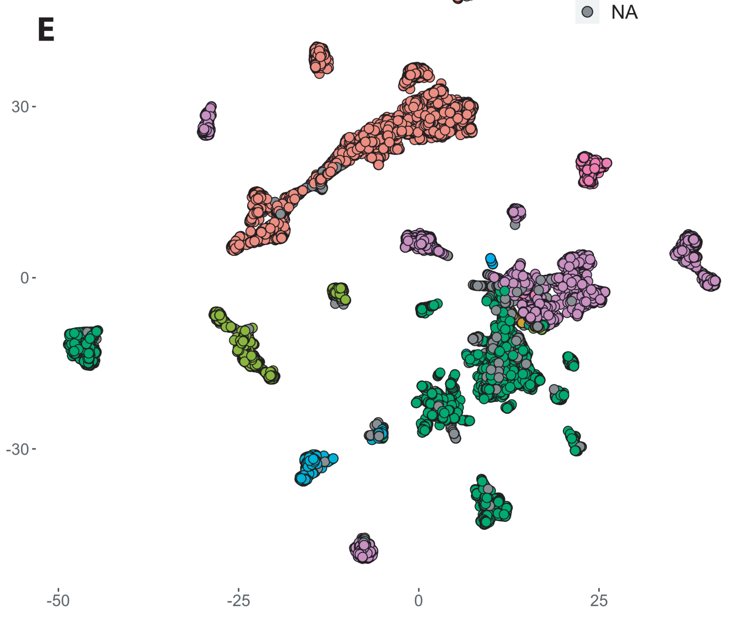


**Supplementary Figure S2**

t-SNE visualizations of ASM-clust output using a matrix generated with MMSeqs2 on five different randomly picked subsets of 1000 sequences (A-E). Although there are small differences between the plots, clustering is robust across all matrices based on five randomly selected reference subsets. One notable exception being the separation of the mannonate dehydratase subgroup in two seperate clusters in E. In panels A, B, and D this separation is visible, but incomplete. The separation in E can be attributed to a high number of the randomly picked reference sequences originating from within the mannonate dehydratase group, providing increased discriminatory power.

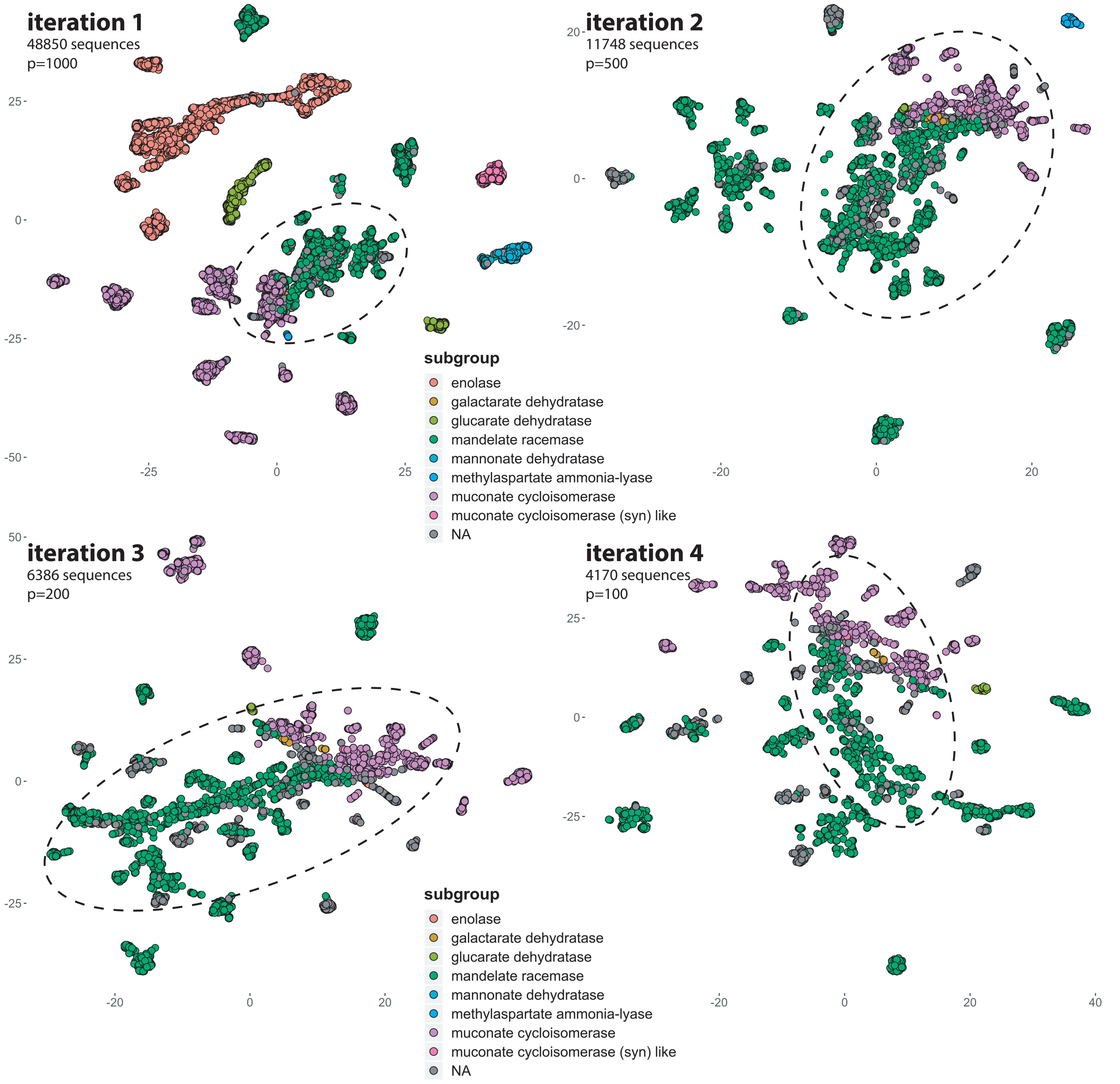


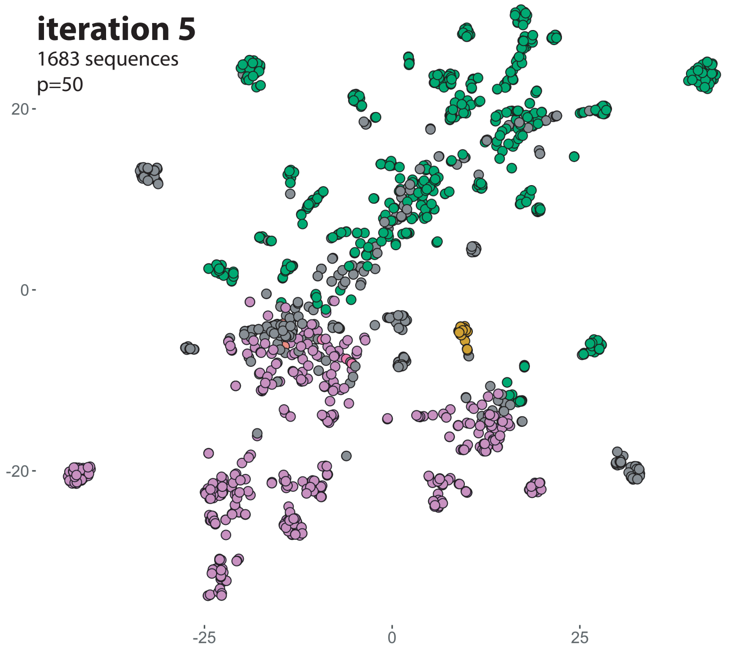


### Supplementary Figure S3

t-SNE visualizations of ASM-clust output on increasingly smaller subsets of the enolase superfamily. Each sequence is displayed as a point, colored by SFLD 'subgroup' annotation. Sequences without SFLD subgroup annotation are colored grey. The number of sequences displayed and the t-SNE perplexity value are indicated on each plot. The dashed circle (roughly) indicates sequences included in the next iteration. The isofunctional Methylaspartate ammonia-lyase (305 sequences) is clearly separated at iteration 2, whereas galactarate dehydratase (25 sequences) is only separated at iteration 5. This can be explained by the perplexity value being set to the same order as the cluster size, and the random subset including a larger number of galactarate dehydratase sequences (due to smaller total sequence size).
